# The temporal dynamics of *same* vs. *different* relational integration in the adult brain

**DOI:** 10.64898/2026.11.26.769422

**Authors:** Zhen Zeng, Antonia Götz, Vishnu Priya Kandasamy, Susan Hespos, Genevieve L. Quek

## Abstract

Humans readily abstract common relations across situations that look entirely different. For example, when reading a map, we align relations between landmarks on a screen with those in the world around us. Relational comparison forms the basis of analogical reasoning. Here, we investigated how recognising *same* and *different* relations instantiated by various simple shapes unfolds in the brain over time. We recorded EEG from adults (*N* = 20) viewing shape pairs presented in a periodic visual oddball paradigm, in which exemplars of one relation occurred at regular intervals within a stream of the other (e.g., *same–same–same–different* or vice versa). Frequency-domain analysis revealed robust responses at the stimulation rate (2.5 Hz) and the oddball rate (0.625 Hz), providing an implicit neural measure of relational discrimination. Time-resolved decoding showed that *same* and *different* could be decoded while holding Sequence Role constant: relation decoding was reliable when both relations served as oddballs, but significantly weaker when they served as standards. *Same/different* decoding at the oddballs generalised to shapes and colours withheld from classifier training and was not explained by stimulus familiarity. These findings show that the brain implicitly registers *same* and *different* relations at an abstract level that generalises across different instantiations of these relations, and that this information is expressed most strongly when a given relation differs from its established context.

**Highlights:**

1. The adult brain registers *same/different* relations implicitly, in the absence of an active analogical reasoning task
2. We isolate relational integration signals that generalise across many unique instances of *same/different* stimuli, minimising the contribution of specific visual features
3. Sensitivity to changes in second-order relations is already evident during perceptual processing, prior to deliberate cognitive evaluation
4. Relational information is encoded through comparison with the surrounding temporal context

## Introduction

Recognising that two situations share a common structure, even when they look nothing alike, is central to human reasoning. When reading a map, for example, we align the spatial relations between symbols on a screen with those between real world locations and landmarks. This ability to abstract a relation from the entities that instantiate it and carry it to new situations, underlies analogical capabilities and abstract thought (Gentner, 1983; 2010). In its simplest form, relational reasoning enables us to perceive commonalities in instances of *same* or *different*: For example, recognising that circle-circle is alike to cross-cross requires setting aside the identity of the objects themselves, and attending to how they relate to one another.

What are the neural processes that underlie our ability to integrate *same*/*different* relations across different instantiations? Although a large visual neuroscience literature has sought to characterise sensitivity to high-level relations between individual objects – for example, their configural arrangement (Ackerman & Courtney, 2012; Kaiser et al., 2019; Quek & Peelen, 2020) or semantic and contextual associations (Faivre et al., 2019; Quek et al., 2025, Yeh et al., 2026) – neural signatures of the relational abstraction involved in *same/different* comparisons are more complex to index. To do so requires operationalising the perception of what Premack (1983) termed **second-order relations** – perceiving that AA is like BB is like CC, even though A, B, and C are distinct – which hold not between individual objects, but between the relations themselves.

Recognising that pairs of objects share a *same* or a *different* relation, irrespective of the objects themselves, has been studied extensively in various species and developmental stages via a behavioural framework (Anderson et al., 2018; Carstensen et al. 2019; Christie & Gentner, 2014; Ferry et al., 2015; Premack, 1983; Smirnova et al. 2015; Wasserman & Young, 2010). Non-human animals can learn to discriminate *same* from *different* pairs, but generalising the relation to novel stimuli often requires extensive training, with variable performance across species and tasks (Wasserman & Young, 2010). In contrast, humans appear to readily abstract relations from a very early age: after brief exposure to training pairs depicting one relation (either *same* or *different*), infants look longer at the previously unseen relation than they do at the learned relation instantiated with novel stimuli (Ferry et al., 2015; Anderson et al., 2018). Generalising relations between sets of different exemplars becomes increasingly flexible across childhood and is facilitated by relational language (Christie & Gentner, 2014; Shivaram, Shao et al., 2026; Shivaram, Götz, et al.,2027). Structure-mapping theory describes how a process of comparison might drive this learning: aligning two pairs of stimuli highlights the relations they share and makes differences within that shared structure especially salient (Markman & Gentner, 1993; Kotovsky & Gentner, 1996; Gentner, 2010). Neuroimaging work has begun to examine this ability: functional MRI studies of relational reasoning implicate posterior and frontal regions, including the rostrolateral prefrontal cortex (Christoff et al., 2001; Wendelken et al., 2008; Davis et al., 2017; for a review see Holyoak & Monti, 2021), and event-related potential (ERP) studies show that the P300 is sensitive to analogical difficulty (Qiu et al., 2008).

Capturing an unambiguous neural index of *same*/*different* discrimination requires dissociating neural responses to these relations from responses to the objects that instantiate them. Auditory oddball designs have achieved this effectively, capturing mismatch negativity (MMN) responses for tone pairs whose relation (e.g., interval direction) deviates from an established one, even as the absolute pitch of the pairs varies (Saarinen et al., 1992; Paavilainen, 2013). At the same time, however, these designs typically do not explicitly test the extent to which neural responses for *same* and *different* generalise across novel instantiations. Fast Periodic Visual Stimulation (FPVS) provides an ideal way to directly address this gap. Partly inspired by vMMN paradigms, FPVS designs embed various exemplars of one category (e.g., faces) at strict periodic intervals within a rapid stream of another (e.g., objects). This stimulation evokes a periodic neural response that is measurable at the scalp level using EEG, with the neural response arising at the embedded frequency providing an objective index of the brain’s differential response to the ‘oddballs’ amongst the ‘standards’ (e.g., Liu-Shuang et al., 2014; Rossion, 2014; Quek & Rossion, 2017; Rossion et al., 2015). Importantly, this oddball response depends critically both on neural responses evoked by the two categories being sufficiently different from one another (i.e., discrimination), as well as on responses within a category being sufficiently similar (i.e., generalisation).

Building on studies that have used FPVS to isolate neural processing of objects’ real-world relations (Quek & Peelen, 2020; Vettori et al., 2026), here we used the approach to obtain an objective neural correlate of *same*/*different* relational processing in the adult brain. We recorded EEG as observers viewed a 2.5 Hz stream of various geometric shape pairs depicting a standard relation, with shape pairs depicting an oddball relation embedded as every fourth stimulus (i.e., SSSD or DDDS). Since standards and oddballs in this design comprise the same constituent elements (re-combinations of 10 unique shape identities), the differential neural response arising at the oddball frequency serves to capture second-order integration of the abstract relations instantiated by the elements within each pair (e.g., *AB* appearing after *AA-BB-CC*, or else *CC* appearing after *AB-CD-EF*). We combined this frequency-based approach with time-resolved neural decoding of both Relation Type (*same* or *different*) and Sequence Role (standard or oddball), enabling us to effectively dissociate neural encoding of a pair’s depicted relation from its status within the sequence. This enabled us to examine evidence for mandatory encoding of *same* and *different* relations (i.e., arising in response to any encounter with the relation), as well as evidence that encoding of a given relation is modulated by its comparative context established by the preceding stimuli. If the brain mandatorily encodes *same* and *different* whenever they are encountered, Relation Type should be equally decodable regardless of whether the stimuli functioned as standards or oddballs. Alternatively, if relational information is brought out by comparison with surrounding context (as structure-mapping theory suggests) *same* and *different* will be most distinguishable when they function as oddballs, where their alignable difference with the surrounding context should strengthen encoding of the relation.

## Method

### Participants

Data from 20 healthy adults (12 females, mean age 29.4 years, SD = 6.67) aged 24-39 were included in the study. Participants were recruited from the undergraduate student population and the community in a major metropolitan Australian city. Two other participants were tested but excluded from the final sample: one due to a history of neurological conditions and the other due to technical error. This sample size is comparable to previous FPVS studies (Vettori et al., 2026). All participants reported having normal or corrected-to-normal vision and no neurological or psychiatric disorders. Participants provided verbal and written consent to complete an hour-long visual EEG task and were compensated $30 AUD. This study has been approved by the Human Research Ethics Committee at 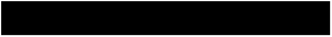 (Ethics No: 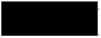).

### Stimuli

Stimuli were constructed from 10 unique geometric shapes (square, circle, hexagon, cross, triangle, trapezoid, semicircle, L-shape, Pac-Man, and raindrop). As shown in Figure 1, each stimulus consisted of a pair of shapes, with *same* pairs containing two of the same shape (e.g., purple L-shape + purple L-shape) and *different* pairs containing two distinct shapes (e.g., grey circle + grey cross). In both cases, shapes within a pair always spanned an identical vertical height and were an identical colour.

**Figure 1.**
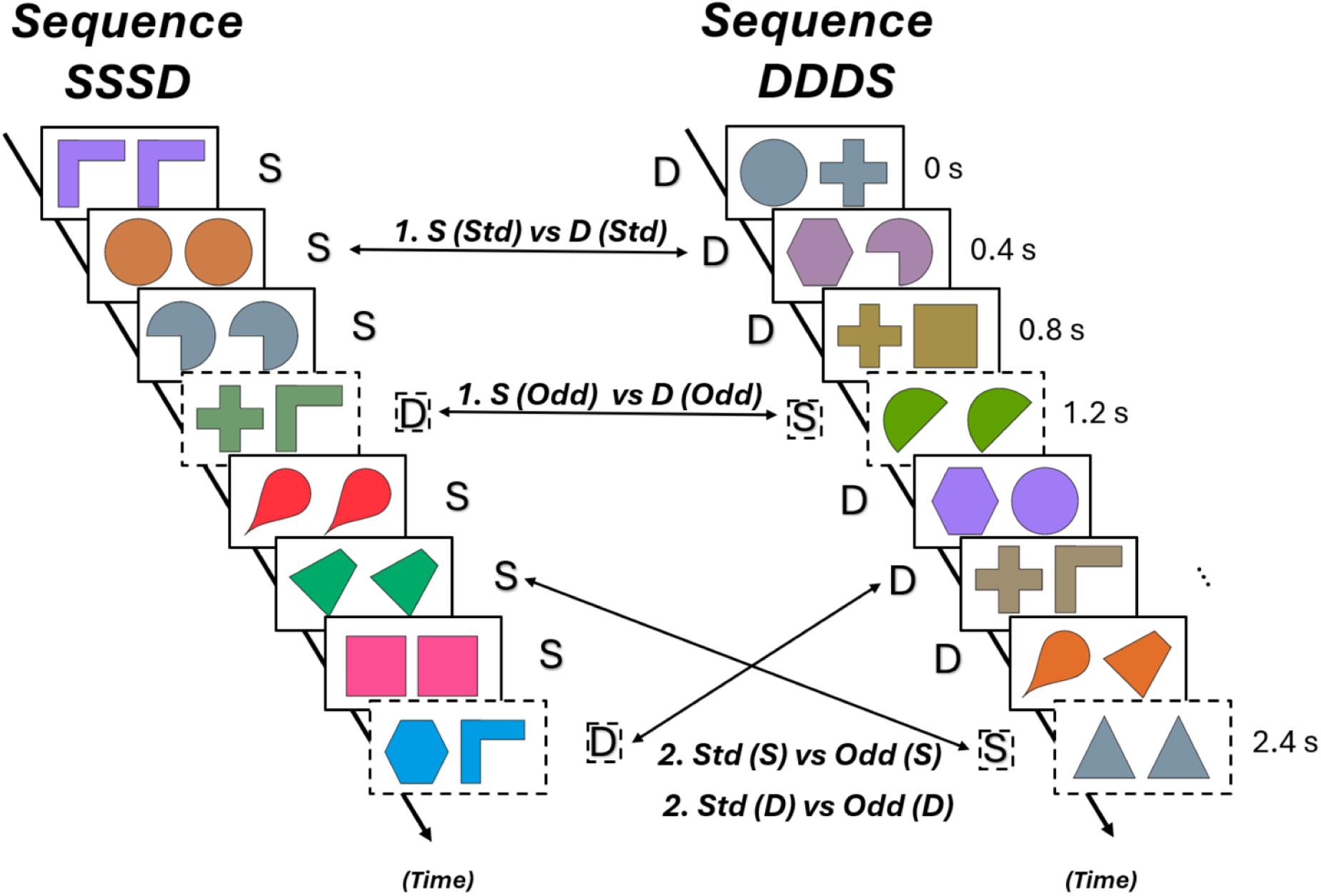
Schematic example of a stimulus sequence and experimental design. In SSSD sequences (at left), ‘standard’ stimuli depicting the *same* relation (solid boxes) appeared at a strict rate of 2.5 Hz, with ‘oddball’ stimuli depicting the *different* relation (dashed boxes) embedded as every fourth stimulus (2.5 Hz/4 = 0.625 Hz). These roles were reversed in DDDS sequences (at right), where *same* and *different* relations served as oddballs and standards respectively. Each stimulus lasted 300 ms and was followed by a 100 ms blank ISI. Note that the box frames surrounding the stimuli are included here to delineate standards/oddballs; these did not appear during the real experiment. Coloured arrows indicate various multivariate decoding analyses enabled by this design: (1) decoding Relation Type with Sequence Role held constant and (2) decoding Sequence Role with Relation Type held constant.

Individual shapes were created in Adobe Illustrator with a 4-pt outline and subsequently assembled into pairs using MATLAB R2025b, with their positions standardised within two adjacent 300 × 300-pixel regions separated by a 50-pixel gap. Stimuli were rendered using an 18-colour approximately isoluminant palette generated in CIE *Lab* colour space in MATLAB, with *L*\* held constant and hue varied systematically by manipulating the *a*\* and *b*\* coordinates at a fixed chroma. The resulting colours were converted to sRGB and applied consistently across all shape stimuli; this served to equate perceived brightness and saturation across colours, so that luminance differences could not influence the frequency-tagging response, attract gaze, or provide an extra cue for relational matching. All final stimuli were exported as PNG images with transparent backgrounds.

The full set of stimuli consisted of 180 *same* pairs and 1620 *different* pairs. The subset of *same* pairs comprised two identical shapes from the 10 shapes listed above rendered in 18 different colours. The subset of *different* pairs comprised non-identical pair combinations of the 10 shapes listed above (45 possible shape combinations; *2 to include reversed positions e.g., circle-square and square-circle = 90) rendered in 18 different colours (90 * 18 = 1620).

During the experiment, the stimulus pairs appeared centrally on a uniform grey background, sized at 400 × 200 px (randomly scaled to 80–120% on each presentation) such that, at a viewing distance of 70 cm, they spanned approximately 9.0° × 4.5° of visual angle at 100% size, ranging from 7.2° × 3.6° (80%) to 10.8° × 5.4° (120%). There was a central fixation cross that turned into a dot during the experimental session (described in **Procedure**). All stimuli are available on osf (xxx).

### Procedure

The experiment was carried out in a quiet, dimly lit testing room that the experimenter visually monitored via webcam from the adjacent room. The design is shown in Figure 1. Participants viewed a continuous stream of object pairs where the standard relation appeared at a rate of 2.5 Hz and the oddball relation was embedded as every fourth stimulus (i.e., 2.5 Hz/4 = 0.625 Hz). Each stimulus was on the screen for 300 ms, followed by a 100 ms inter-stimulus interval. Each trial comprised four stimuli: three standards followed by an oddball (*same*–*same*–*same*–*different* in SSSD sequences; *different*–*different*–*different*–*same* in DDDS sequences); this meant that the oddball presentation rate was 2.5 Hz / 4 = 0.625 Hz). Within each sequence, stimuli were sampled at random without repetition, and consecutive stimuli never repeated a shape or colour. As such, the oddball was discernible from surrounding standards by virtue of the relation between the two shapes in the pair, rather than any specific visual features.

Each sequence contained 37 trials (37 trials × 4 stimuli × 400 ms = 59.2 s), with the first and last two trials being gradually faded in/out respectively in terms of maximum stimulus contrast (these four trials were excluded from analysis). Thus, the sequence proper contained 33 full contrast trials and lasted 52.8 seconds. Each participant completed 20 sequences (10 SSSD and 10 DDDS), organised into 10 reshuffles, each comprising one SSSD and one DDDS sequence in random order; sequences were separated by self-paced breaks. Across the full experiment, each participant saw 1480 instances of *same* and 1480 instances of *different* (1320 each excluding the fade-in/out trials). This meant that within the experiment, the participant saw all 180 unique *same* stimuli and approximately 970 of the 1620 unique *different* stimuli. In other words, over the 10 reshuffles, *same* stimuli were necessarily reused ∼8 times and *different* stimuli ∼1.5 times.

To maintain vigilance, participants performed an orthogonal fixation task, as is common in rapid visual presentation EEG studies (Grootswagers et al., 2022, 2024; Huang et al., 2026; Shatek et al., 2022): they pressed the spacebar whenever the central fixation cross changed into a dot (2–5 times per sequence). This task performance was not analysed further.

### EEG Acquisition

We recorded scalp EEG using a 64-channel Biosemi system (standard 10-10 system) at a sampling rate of 1000 Hz. Impedance was reduced below 25 kOhm at each electrode site whenever possible using conductive gel. All stimuli were presented using a customised script run in Psychopy3 onto a Viewpixx 120 Hz monitor. We synchronised the EEG recording at the start of the sequence as well as on each stimulus by using a viewpix pixel trigger system to send triggers to the EEG acquisition PC.

### Analysis

#### Preprocessing

Each participant’s continuous EEG signal was downsampled to 250 Hz, band-pass filtered (0.1–100 Hz) and notch filtered (49–51 Hz band-stop), all with fourth-order Butterworth filters, and re-referenced to the average of all scalp channels. Minimum preprocessing was performed with FieldTrip (version 20201229; Oostenveld et al., 2011) in MATLAB R2023b. For the frequency-domain analysis only, ocular artefacts were additionally removed with independent component analysis before re-referencing (see Frequency-Domain Analysis); the decoding analyses used the preprocessed data without this step. Epoching procedures specific to each of the two analyses are described in the corresponding sections below. For all analyses, the first two and last two trials of each sequence (fade-in and fade-out) were excluded.

#### Frequency-Domain Analysis

Frequency-based quantification of the periodic responses was used to examine neural discrimination of *same* and *different* relations presented within the same sequence. Because no electro-oculogram was recorded and the oddball frequency (0.625 Hz) lies within the frequency range of blinks, ocular artefacts were removed before this analysis, as in previous FPVS studies (Retter & Rossion, 2016) Independent component analysis (runica, 64 components) was fitted to each participant’s filtered data before average re-referencing, using a 1 Hz high-passed copy of the stimulation sequences. Components reflecting blinks or eye movements were identified by visual inspection of their topographies and time courses, aided by their correlation with frontal electrodes (mean of Fp1 and Fp2; AF7 minus AF8), and were removed from the 0.1 Hz data (1 component for 18 participants, 2 for one participant and none for one participant), after which the data were average referenced. This analysis was performed at the sequence level. Each sequence was segmented from the onset of its third trial (i.e., excluding the two fade-in trials) and cropped to 30 oddball cycles (≈ 47.75 s), using the stimulation rate measured from the triggers (mean stimulus onset asynchrony 397.9 ms), so that the base and oddball frequencies fell exactly on frequency bins (resolution ≈ 0.021 Hz). Sequences were averaged in the time domain separately for each sequence type (SSSD, DDDS) and for both combined, and the amplitude spectrum (µV) of each average was computed with a fast Fourier transform (FFT) for each electrode. Frequencies are reported relative to the nominal stimulation rates (2.5 Hz and 0.625 Hz).

To identify the harmonics that carried signal, we followed Quek and Peelen (2020): the amplitude spectra of both sequence types combined were averaged across participants and then across all 64 channels to produce a single grand-average spectrum, so that harmonic selection was blind to Sequence Type. The amplitude at each frequency bin was converted to a z-score relative to a local noise range of the 8 surrounding bins on each side, excluding the immediately adjacent bin, any bins falling on other oddball or base-rate harmonics, and the single largest and single smallest amplitude among the remaining bins. Oddball harmonics (multiples of 0.625 Hz that are not multiples of the base rate, inspected up to 30 Hz) and base-rate harmonics (multiples of 2.5 Hz, inspected up to 47.5 Hz) were considered significant if z > 1.64 (p < .05, one-tailed), inspecting harmonics in ascending order until two consecutive harmonics were not significant. This identified 11 oddball harmonics (0.625– 8.75 Hz) and 17 base-rate harmonics (2.5–45 Hz, all except 35 Hz). Baseline-corrected amplitude spectra (SBL) were computed at every frequency bin, channel, participant and sequence type by subtracting the mean amplitude of the same local noise range from the amplitude at that bin. The summed oddball response was quantified as the sum of SBL over the significant oddball harmonics, and the summed base response as the sum over the significant base-rate harmonics, separately for each channel and sequence type. There is no established topography for a response to a relational change; to avoid selecting a region of interest from the same data, summed SBL was averaged across all 64 channels, and topographies are shown for description only. Whole-scalp summed SBL was tested against zero with one-tailed one-sample *t*-tests and JZS Bayes factors (Cauchy prior scale √2/2), separately for SSSD and DDDS sequences, and the two sequence types were compared with paired two-tailed *t*-tests and Bayes factors with the same prior; the summed base response was tested in the same way. At the individual level, a summed response was considered significant when the z-score of the participant’s whole-scalp spectrum, summed over segments centred on the significant harmonics, exceeded 1.64.

#### Time-resolved multivariate decoding analysis

The following decoding analyses were conducted: Contrast 1 tested the neural discriminability of a stimulus’ depicted Relation Type (*same* or *different*), while holding Sequence Role constant. Contrast 2 tested the neural discriminability of Sequence Role (standard or oddball), while holding Relation Type constant (see details below in Decoding Contrasts). Control analyses with strict shape and colour hold-out and further checks were also conducted, these are provided in Supplementary Material. The procedure described in this section applies to all analyses unless stated otherwise.

All decoding analyses were performed at the stimulus level. We segmented a 900 ms epoch time-locked to the onset of each 300 ms stimulus presentation, lasting from −100 to 800 ms. Each resulting epoch was baseline corrected using the pre-stimulus interval (−100 to 0 ms). Multivariate pattern analysis was performed using CoSMoMVPA (Oosterhof et al., 2016) and FieldTrip in MATLAB. Decoding used a leave-one-reshuffle-out cross-validation scheme (a reshuffle being one SSSD and one DDDS sequence; see Procedure): in each fold, a linear discriminant analysis (LDA) classifier was trained to discriminate the relevant classes for that specific contrast using the epochs from all reshuffles except one and tested on epochs from the held-out reshuffle. This was repeated until every reshuffle had served as the test set exactly once, with classification accuracy averaged across folds. Within each fold, training-set classes were balanced by random subsampling to the size of the minority class. Spatiotemporal EEG patterns across all electrodes were used as features. Decoding accuracy was computed across a sliding window of 20 ms width, stepped in 12 ms increments across the epoch, yielding a time series of classification accuracy per participant. Group-level statistical inference used Bayesian one-sample *t*-tests against chance (0.5) at each time point, implemented via the BayesFactor R package with a medium prior scale. Bayes Factors (BF_10_) are reported for each time point, with BF_10_ ≥ 3 treated as moderate evidence for above-chance decoding.

Spatial analysis used a channel searchlight approach in which the decoder described above was applied separately for each electrode and its four nearest neighbours (as defined by the biosemi64 layout), producing a Bayes Factor map across the scalp at each time point. Group-average searchlight results were visualised as topographic maps at 100 ms intervals from 100 to 600 ms post-stimulus onset.

To assess the temporal stability and generalisation of neural representations, a temporal generalisation matrix (TGM; King & Dehaene, 2014) was constructed by training the classifier on each time window and testing it on every other time window; its diagonal corresponds to the sliding-window result. Significant regions of the TGM were identified with a cluster-based permutation test in two steps. First, clusters were identified: at every cell, participants’ accuracies were tested against chance (0.5) with a one-sample *t*-test, cells with above-chance accuracy (a positive t-value exceeding the two-tailed p < .05 threshold) were retained, retained cells that were adjacent in training or testing time were grouped into clusters, and each cluster’s mass was defined as the sum of its t-values. Because only above-chance cells could form clusters, only above-chance generalisation was tested. Second, the clusters were tested against a null distribution of cluster mass built from 1,000 sign-flip permutations: in each, every participant’s accuracy-minus-chance matrix was multiplied by +1 or −1 (one sign per participant, preserving the temporal autocorrelation of the TGM), clusters were identified again in the same way, and the largest cluster mass was retained (0 if no cluster formed). Observed clusters whose mass exceeded the 95th percentile of this distribution were considered significant (one-tailed, family-wise α = .05). The procedure was run separately for each decoding contrast.

Two sets of paired contrasts were used to examine how Relation Type (*same* or *different)* and Sequence Role (standard or oddball) contributed to the EEG signal (summarised in Figure 1). As each stimulus presentation lasts 300 ms (plus 100 ms ISI), the standard class in our decoding analyses was always represented by the neural response evoked by the second standard in the trial; this ensured no overlap between the standard and oddball epochs.

##### Contrast 1: Decoding Relation Type

The first set of contrasts asked whether neural responses to *same* and *different* pairs were discriminable when Sequence Role is held constant. To this end, we decoded the second standard of S**S**SD from the second standard D**D**DS (i.e., both standards), and, separately, the oddball of SSS**D** against the oddball of DDD**S** (i.e., both oddballs). Above-chance decoding here would suggest the neural response contains information about the stimulus’ Relation Type (*same* or *different*) that cannot be accounted for by its role within the sequence.

##### Contrast 2: Decoding Sequence Role

Our second set of contrasts asked whether neural responses to standards and oddballs are discriminable when Relation Type is held constant. To this end, we discriminated responses evoked by *same* pairs functioning as the second standard (sequence SSSD) against *same* pairs functioning as the oddball (sequence DDDS) and then repeated the equivalent analysis for *different* pairs. Because the two stimulus types being classified share the same Relation Type but differ in role and originate from different sequences, above-chance decoding indicates that the neural response to a stimulus depends on its Sequence Role when its own Relation Type is held constant. Because Sequence Role is defined by the surrounding stimuli, which differ in Relation Type between the two sequences, this decoding can also include responses to neighbouring stimuli that overlap the epoch.

###### Stimulus and Artefact Controls

To test whether *same/different* decoding could be explained by the same shapes or colours appearing in both the training and test sets, Contrast 1 analyses (decoding Relation Type) were also conducted using strict controls in which either one stimulus shape or one stimulus colour was held out; Contrast 2 analyses (decoding Sequence Role) were likewise repeated with the shape hold-out. In this analysis, the test stimuli shared no shape (colour) with any training stimulus, so above-chance decoding required generalisation to unseen shapes and colours (Supplementary Material S1). Further controls are described in the Supplementary Material: (1) matching *same* and *different* pairs on prior exposure and comparing early with late reshuffles **(S2)**, (2) normalising each presentation’s activity pattern for amplitude, to test whether decoding reflected differences in overall response size **(S3)**, and (3) constraining decoding to use only the 20 posterior channels re-referenced to their own mean, thereby preventing ocular activity from contaminating the signals via re-referencing **(S4)**.

## Results

### Frequency-Domain Results

The grand-average baseline-corrected spectrum showed clear peaks at the base rate (2.5 Hz) and its harmonics, and at the oddball rate (0.625 Hz) and its harmonics (Figure 2B). The summed oddball response (11 harmonics, 0.625–8.75 Hz) was reliably above zero in both SSSD sequences (M = 0.31 µV, SD = 0.27; t(19) = 5.11, p < .001, BF₁₀ = 428) and DDDS sequences (M = 0.45 µV, SD = 0.36; t(19) = 5.66, p < .001, BF₁₀ = 1,260), confirming that the brain discriminated oddballs from standards whether the oddball was a *different* or a *same* pair. Because shapes and colours changed on every stimulus, an oddball response requires the brain to treat the oddball differently from the preceding standards on the basis of the relation between its two shapes; it cannot, however, separate the relation itself from its role in the sequence, which the decoding contrasts address. The summed oddball response was larger in DDDS than in SSSD sequences (difference = 0.15 µV, SD = 0.19; t(19) = 3.46, p = .003, BF₁₀ = 16) and was largest over occipito-parietal electrodes in both sequence types (Figure 2C). At the individual level, 14 of the 20 participants showed a significant summed oddball response. Without ICA, the summed oddball response was also above zero in both sequence types (SSSD: BF₁₀ = 29; DDDS: BF₁₀ = 23), but the difference between them was inconclusive (BF₁₀ = 0.56). The summed base response (17 harmonics, 2.5–45 Hz), which reflects general visual stimulation and is not of theoretical interest here, was strong in both sequence types (SSSD: M = 1.37 µV, t(19) = 11.87, BF₁₀ = 3.2 × 10⁷; DDDS: M = 1.36 µV, t(19) = 12.22, BF₁₀ = 5.0 × 10⁷), significant in all 20 participants, maximal over medial occipital electrodes, and did not differ between sequence types (t(19) = 0.17, p = .87, BF₀₁ = 4.3).

**Figure 2.**
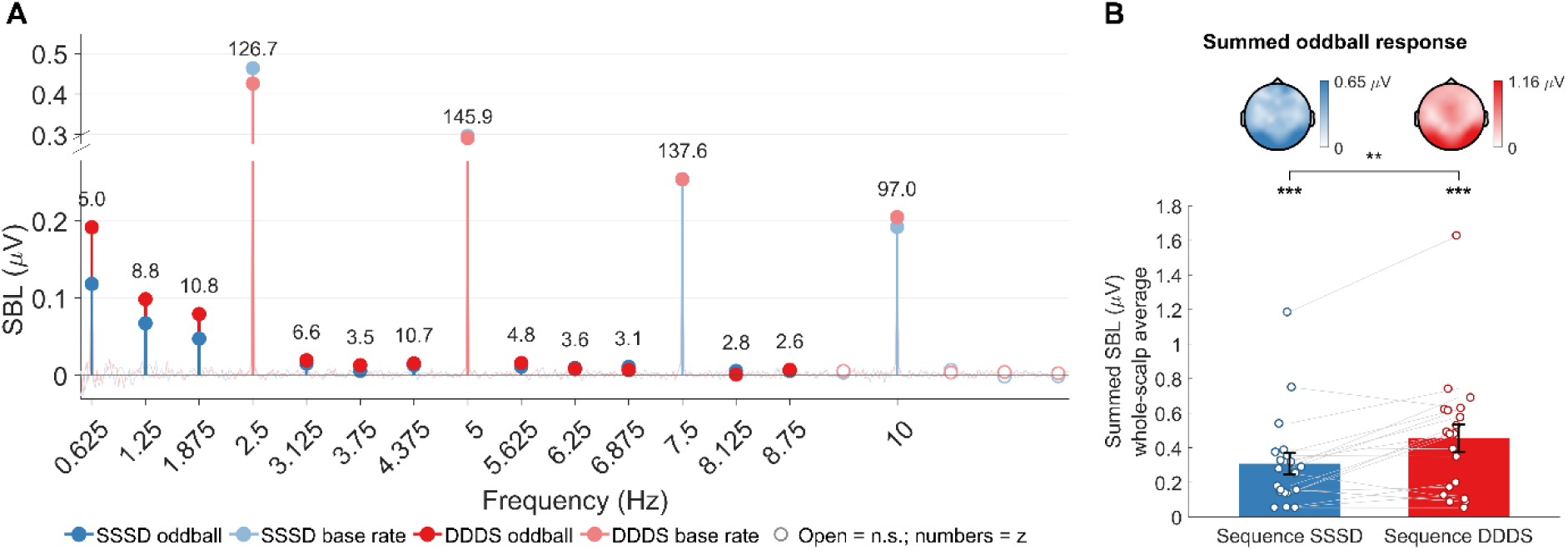
Frequency-domain responses. **A**. Grand-average baseline-corrected amplitude spectrum for SSSD (blue) and DDDS (red) sequences (average of all 64 channels), 0.5–12 Hz, shown as stems at the harmonic frequencies over the full spectrum (faint lines). Filled dots mark significant harmonics (oddball harmonics in a lighter shade, oddball-rate harmonics in full colour) and open dots non-significant ones; numbers give the *z*-score of each significant harmonic (both sequence types combined). The y-axis is broken so that the oddball harmonics are visible beneath the much larger stimulation-rate peaks. **B**. Summed oddball response (whole-scalp summed SBL over the 11 significant oddball harmonics) in SSSD and DDDS sequences. Bars show the group mean ± SEM; dots and grey lines show individual participants; topographies above each bar show the group-mean summed SBL at each channel (shared colour scale). Asterisks above the bars: summed SBL > 0 (one-tailed *t*-test); bracket: SSSD vs. DDDS (paired *t*-test); *** p < .001, ** p < .01, n.s. not significant.

### Multivariate Decoding Results

#### Contrast 1: Decoding Relation Type

To investigate how neural responses encode information about *same* and *different* relations independently of encoding their Sequence Role, we trained a classifier to discriminate these classes exclusively based on epochs corresponding to the second standard in the trials, or to the oddballs in the trials (see Methods). Figure 3A shows that participants’ neural signals contained decodable information about the depicted relation in both cases, although the distinction between *same/different* was notably stronger for the analysis constrained to oddball instances of these relations, showing sustained above-chance decoding between ∼100 and 550 ms (peak accuracy ∼57% at ∼290 ms). When both relations were standards, decoding was much weaker (peak accuracy ∼52% at ∼110 ms) and exceeded the evidence threshold only intermittently across the epoch, without a sustained period of above-chance decoding. A paired Bayesian comparison of the two decoding time courses indicated stronger decoding for oddballs than for standards between ∼150 and 520 ms (grey squares in Figure 3A). *Same* and *different* pairs were thus distinguished from about 100 ms, but the dependence of this distinction on Sequence Role emerged from about 150 ms.

**Figure 3.**
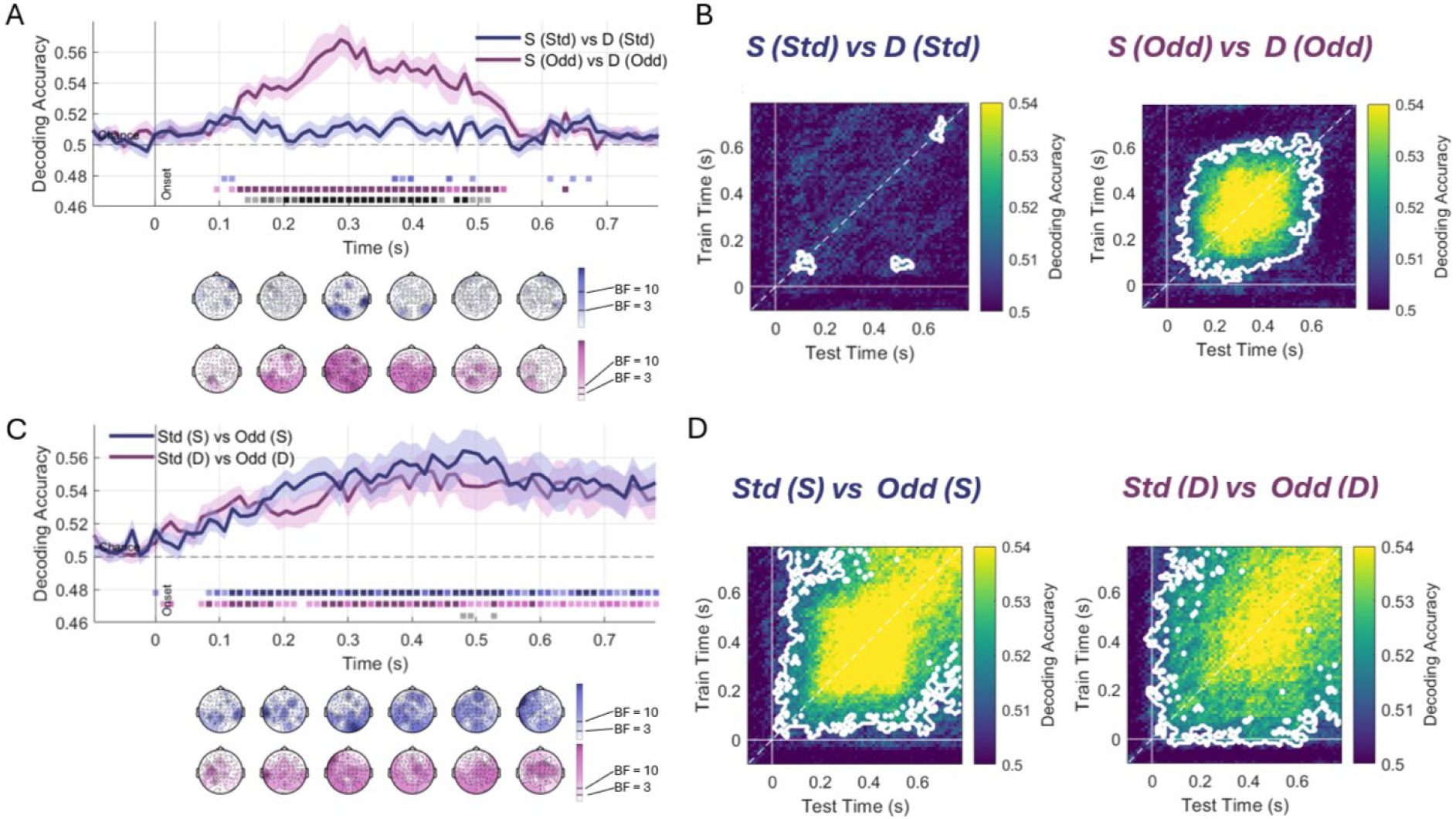
Time-resolved decoding of Relation Type and Sequence Role. (A) Contrast 1, Relation Type: decoding accuracy over time for S (Std) vs. D (Std) (dark blue) and S (Odd) vs. D (Odd) (purple), shown as mean ± SEM; the dashed line marks chance (50%) and the vertical line stimulus onset. Coloured squares below the curves mark time points with evidence for above-chance decoding (BF₁₀ ≥ 3, ≥ 10 and ≥ 30, in increasing colour intensity), and grey squares evidence for a difference between the two curves. The scalp maps beneath show searchlight log₁₀ BF₁₀ from 0.1 to 0.6 s (upper row, S (Std) vs. D (Std); lower row, S (Odd) vs. D (Odd)); only channels with BF₁₀ ≥ 3 are coloured. (B) Temporal generalisation matrices for the two contrasts in A, showing decoding accuracy when the classifier was trained at one time (y-axis) and tested at another (x-axis); the dashed line marks the diagonal (train time = test time) and white contours mark significant clusters (cluster-based permutation test, p < .05). (C) Contrast 2, Sequence Role: decoding accuracy for Std (S) vs. Odd (S) (dark blue) and Std (D) vs. Odd (D) (purple), with searchlight maps, plotted as in A. (D) Temporal generalisation matrices for the two contrasts in C, plotted as in B.

Temporal generalisation analysis mirrored these results (Figure 3B): where same/different decoding for oddballs gave rise to a single large cluster, spanning ∼100–550 ms along the diagonal and generalising across training and testing times between ∼0 and ∼650 ms, the same analysis for standards showed only small, isolated clusters (around 100 ms and 650 ms on the diagonal, and one off-diagonal cluster in which classifiers trained at ∼100 ms generalised to ∼500 ms).

#### Contrast 2: Decoding Sequence Role

These contrasts asked whether standards and oddballs were discriminable when Relation Type was held constant. Decoding was above chance for both contrasts from about 100 ms onwards and remained so for most of the epoch (Figure 3C). This did not depend on the particular shapes presented: when the classifier was tested only on pairs whose shapes it had never encountered during training (leave-shapes-out cross-validation), Sequence Role was still decoded above chance, at a level close to that of the original cross-validation (200–600 ms: Std (S) vs. Odd (S) 55.2%, BF₁₀ = 170, versus 55.4%; Std (D) vs. Odd (D) 53.4%, BF₁₀ = 18, versus 54.3%; Supplementary Material S1). This indicates that the response to a stimulus depends on its Sequence Role when its own Relation Type is held constant. Searchlight maps showed that this information was broadly distributed across the scalp from 100 ms onwards. Paired Bayes factors showed no reliable difference in decoding accuracy between the two contrasts: evidence for a difference reached BF₁₀ ≥ 3 at only two time points (around 480–500 ms), which we do not interpret. Temporal generalisation (Figure 3D) yielded a single large cluster for each contrast that extended across most of the epoch, indicating that the information distinguishing standards from oddballs was sustained and generalised broadly across time.

### Stimulus and Artefact Controls

To confirm that *same/different* decoding could not be accounted for by shape- or colour-specific factors, we repeated Contrast 1 with the test pairs restricted to shapes or colours absent from training. S (Odd) vs. D (Odd) decoding remained well above chance (leave-shapes-out: 53.9%; leave-colours-out: 54.4%; both BF₁₀ > 10,000), as did S (Std) vs. D (Std) (BF₁₀ = 3.8–67; Supplementary Material S1). To confirm that the unequal exposure of individual *same* and *different* pairs (each same pair was shown more often) could not account for the results, we matched *same* and *different* oddballs on prior exposure: decoding remained above chance (52.6%, BF₁₀ = 43), did not differ from a reference with the same number of stimulus presentations (BF₀₁ = 2.8), and did not increase from early to late reshuffles (Supplementary Material S2). To test whether *same* and *different* oddballs differed only in overall response size, we normalised each presentation’s activity pattern for amplitude: decoding was almost unchanged (S (Odd) vs. D (Odd), 120–400 ms: 54.2% versus 54.9%, BF₁₀ = 7,676; Supplementary Material S3), so the two kinds of oddball differed in the shape of their scalp pattern rather than only in its overall size. Finally, to rule out a contribution of eye movements and blinks, we decoded from posterior channels re-referenced to their own mean, which reproduced every contrast with the same size and time course (S (Odd) vs. D (Odd), 120–400 ms: 55.6% versus 54.9% from all channels; Supplementary Material S4).

## Discussion

The present study aimed to provide an unambiguous neural index of *same*/*different* relational encoding in the adult brain, taking advantage of a visual oddball paradigm in which we evoked the standard/oddball distinction in relational properties alone. We presented observers with *same* and *different* shape pairs comprised of the same constituent shape elements (SSSD or DDDS), such that the oddball was identifiable as distinct from the preceding standards only in terms of the relation depicted by the two elements. Frequency analysis revealed a robust response to periodic changes in Relation Type, and multivariate decoding showed that the EEG signal carried information about both Relation Type (*same* vs. *different*) and Sequence Role (standard vs. oddball) independently. Information about *same/different* relations was strongest in neural responses evoked by oddballs (i.e., when that relation violated the regularity established by the sequence context). Together, the findings suggest that relational information becomes available rapidly and generalises across exemplars.

### Frequency tagging reveals relational discrimination across changing exemplars

The frequency-domain response measured here provides some of the first neural evidence that the brain reliably discriminates *same* from *different* relations, even as the exemplars instantiating these relations changed. A reliable response emerged at the oddball frequency representing relational change, suggesting that seeing a relation which deviated from those preceding it drove a differential neural response that was measurable at the scalp level. This effect was present both for *same* oddballs embedded in a stream of *different* standards, and vice versa; slightly stronger in the former condition. Important to note here is that this signal is not attributable to periodic repetition of a specific visual stimulus but rather indexes the visual system’s reliable discrimination of changes in relational content (Quek & Peelen, 2020). This extends FPVS work showing categorical discrimination across faces (Liu-Shuang et al., 2014; Rossion et al., 2015), objects (Rostami et al., 2026) and spatial relations (Vettori et al., 2026), as well as work using FPVS to index high-level relations between real-world objects (Quek & Peelen, 2020), to an abstract relation between simple shapes. Crucially, an oddball response of this kind can arise only if the responses to different instances of a relation are sufficiently similar to one another, and at the same time sufficiently different from the responses to the standards. Because the pairs representing the two relations spanned a wide range of shape and colour combinations (see Figure 1), any oddball response they evoke necessarily reflects a discrimination of *same* and *different* that generalises across highly variable stimuli instantiating these relations. Together these findings provide evidence that this logic can be used to index a high-level, abstract relation such as *same*/*different*.

### *Same* and *different* are best distinguished at the oddball

Turning from frequency-tagging to time-resolved neural decoding, here we found evidence to suggest the brain’s ability it discern between *same* and *different* depended on the relative importance of the stimulus within the sequence: Neural responses evoked by *same* and *different* pairs were distinguishable from ∼100 to 550 ms, but only when each relation broke the regularity of the preceding sequence (i.e., when they functioned as oddballs). When the same exact stimuli functioned as standards – and therefore repeated the preceding regularity – the distinction between *same/different* was much weaker, reaching the evidence threshold only intermittently and without a sustained period of above-chance decoding. Together, these results suggest that relational information is expressed or encoded most strongly when the depicted relation deviates from the context that has been established by the preceding sequence of stimuli. Importantly, weaker decoding of *same/different* relations depicted by standards does not imply that their relation went unrepresented: Indeed, the presence of a measurable oddball response presupposes this, because an oddball is defined only with respect to the relation that the standards share. Rather, we interpret this finding in the context of repetition suppression accounts (Grotheer & Kovács, 2016; Feuerriegel et al., 2018): a relation that repeats is predictable and evokes a weaker, less separable response than a relation that does not repeat, as at the oddball.

Notably, the neural separability of *same* and *different* relations depicted by oddballs was not accounted for by a mere difference in amplitude of the evoked response: *same/different* decoding survived amplitude normalisation (Supplementary Material S3), suggesting the two kinds of oddball evoked qualitatively differing scalp patterns discernible to the linear classifier. Prior work suggests that prediction and surprise alter the information carried by sensory responses, not only their amplitude (Kok et al., 2012; Tang et al., 2018). Indeed, the conceptually-related MMN response has been shown to track violations of abstract regularities, such as the direction of pitch change within tone pairs whose absolute pitch varies (Saarinen et al., 1992; Paavilainen, 2013), or rules defined over whole sequences (Bekinschtein et al., 2009). The present results extend this to the visual modality, and more specifically to the abstract relation between object pairs whose features change on every presentation. Even though the relations were task-irrelevant, participants may have implicitly tracked the oddball regularity as a rule, and the rule a participant is applying is itself encoded in patterns of brain activity and can be decoded from them (Woolgar et al., 2011, 2015). The novel contribution here is that a relation between two objects, instantiated by shapes and colours that changed between consecutive stimuli, can be decoded from time-resolved neural activity.

### Sequence Role tracks the relation, not specific stimuli

Whether a stimulus was a standard or an oddball (Sequence Role) was decodable from about 100 ms after stimulus onset and for most of the epoch, independent of the allocation of the *same* or *different* relations to the standards and oddball roles (Contrast 2). Standards repeat the current regularity, and oddballs break it. In visual oddball paradigms, these roles differ through at least two mechanisms: repetition-related changes in the sensory response to standards, and comparison of the oddball with the regularity set up by the standards (Kimura et al., 2009). In fast periodic oddball designs specifically, immediate stimulus repetition can abolish expectation effects (Feuerriegel et al., 2018), and the mismatch responses behave as all-or-none surprise signals rather than graded expectation suppression (Feuerriegel et al., 2021). Here, consecutive stimuli never shared a shape or colour, so stimulus-specific repetition cannot account for the Sequence Role effect: what repeats from one standard to another is the nature of the relation depicted by the shape pairs. In this way, above-chance decoding of Sequence Role captures the brain’s encoding of relational regularity (established by standards) and relational violations (enacted by the oddball).

### Relational information emerges through comparison

In the present data, neural encoding of *same/different* relations was strongest when that relation violated the relational regularity established by the preceding stimuli, and comparatively weaker when it repeated the preceding relation. This connects to Premack’s (1983) distinction between first-order relations (judging whether two objects are identical) and second-order relations (judging whether the relation between one pair matches the relation between another pair). The present paradigm taps the second-order relation, with an oddball response inherently reflecting the brain’s detection of a mismatch between the relation depicted by the oddball and the ones instantiated by the preceding standards.

Structure-mapping theory provides a framework for why expression of relational information depends on sequence role (Gentner, 1983; 2010), via the process of <u>comparison.</u> Under this rationale, the relation depicted in each standard is ‘aligned’ with that contained in the preceding ones. Since successive standards share a relation, this alignment yields a commonality that helps the viewer abstract the relation from the specific shapes (Kotovsky & Gentner, 1996; Gentner, 2010). In contrast, comparison of an oddball with the preceding standards yields an alignable difference in the depicted relations, such that the oddball’s relation becomes especially salient (Markman & Gentner, 1993; Gentner & Markman, 1997). Evidence in support of this account has come mainly from behavioural tasks; the present study provides implicit neural evidence consistent with the idea that alignable differences are where relational information is most strongly expressed.

### Relational mismatch is registered early and implicitly

The time course we observe here suggests that this signal of relational integration arises relatively early during visual processing. *Same* and *different* pairs were distinguished from about 100 ms, and the dependence of this distinction on Sequence Role, the signature of a mismatch between relations, emerged from about 150 ms. An effect this early invites the question of whether it reflects low-level visual differences between the stimuli rather than the relation between the two shapes, and several aspects of the design and analyses speak against this. First, shapes and colours changed on every presentation, and consecutive stimuli never shared a shape or a colour, so no individual visual feature repeated at the oddball rate. Second, decoding generalised to pairs whose shapes or colour the classifier had never encountered during training (Supplementary Material S1), so it could not rely on the specific shape or colour features. Third, decoding was almost unchanged when the overall amplitude of each activity pattern was normalised (Supplementary Material S3): *same* and *different* oddballs evoked differently distributed patterns of activity, not merely responses of different strength. Fourth, the effects were reproduced from posterior channels re-referenced to their own mean (Supplementary Material S4), so they were unlikely to be carried by eye movements or blinks. Finally, the latency of the mismatch effect falls in the range of mismatch responses to abstract regularities (Saarinen et al., 1992; Paavilainen, 2013) and of visual mismatch responses (Kimura et al., 2009; Smout et al., 2019). Relational mismatch is thus registered rapidly and implicitly over posterior sites, well before the deliberative processes associated with analogical reasoning (Christoff et al., 2001; Davis et al., 2017). Such signals may provide the input on which explicit relational comparison builds, rather than constituting analogical reasoning themselves.

### Limitations and future directions

While these data provide an existence proof that FPVS can be applied to relational discrimination across exemplars, there are some limitations that can be improved upon in future studies. First, *same* pairs consisted of two identical shapes, so the *same* relation cannot be fully separated from the perceptual repetition of identical forms. Future work could separate the two by defining *same* and *different* on another dimension, such as size, using pairs of non-identical shapes that are the *same* or *different* in size, and by testing whether a classifier trained on one dimension generalises to the other within participants, which would show a relational code shared across dimensions. Second, participants viewed the sequences passively, so the present data establish implicit neural discrimination of relations but do not link it to explicit relational judgements. Future studies could combine the present paradigm with an explicit relational matching task, to test whether the strength of the neural response to relational oddballs predicts the accuracy or speed of relational judgements in the same participants.

Finally, because the paradigm requires no explicit response, it is particularly well suited to populations whose behavioural responses are difficult to measure, such as infants and young children. Behavioural work shows that relational generalisation develops substantially across infancy and childhood: infants can generalise *same/different* relations under appropriate learning conditions (Anderson et al., 2018; Ferry et al., 2015), and relational reasoning becomes increasingly flexible across childhood and is influenced by relational language (Hochmann et al., 2017; Shivaram, Shao et al., 2026). The present approach thus offers a promising way to chart the developmental trajectory of relational reasoning, including when the neural distinction between *same* and *different* first emerges, and when it begins to depend on comparison with the surrounding context.

## Conclusion

The present study shows that EEG responses to *same* and *different* pairs carry information about the relation between the objects and the role of standard or oddball. Frequency tagging demonstrated reliable relational discrimination across changing visual exemplars. In addition, time-resolved decoding revealed distinct temporal profiles for the abstract relational and the role the stimuli played in the sequence. Relational information generalised to new shapes and colours, but was expressed most strongly when a relation violated the established regularity. These findings provide an implicit neural account of how relational processing unfolds over time. They show that the brain discriminates *same* from *different* and generalises each relation across new shapes and colours; whether this reflects a fully abstract representation of the relation, independent of any specific visual dimension, remains to be tested. More broadly, the findings provide evidence that relational information is expressed through comparison with an ongoing relational structure, providing a framework for investigating how representations of relations become sufficiently independent of specific visual instances to support the flexible relational comparisons required for analogical reasoning.

## Acknowledgement

This research was partially supported by the Australian Government through the Australian Research Council’s Discovery Projects funding scheme (project DP250101325) awarded to S. Hespos and G. Quek. The views expressed herein are those of the authors and are not necessarily those of the Australian Government or the ARC.

