## Supplementary Material S1-S4 for "The temporal dynamics of *same* vs. *different* relational integration in the adult brain"

### General procedure

Four control analyses test alternative explanations of the *same/different* decoding reported in the main text. Unless stated otherwise, all used the main decoding pipeline. Stimulus presentations were epoched from −0.1 to 0.8 s and baseline-corrected to the pre-stimulus interval (−0.1 to 0 s); the fade-in and fade-out trials of every sequence were excluded. Features were the voltages of all 64 channels at the five samples (250 Hz) of a 20-ms window, i.e. 320 values per presentation, and windows were advanced in 12-ms steps (three samples). A linear discriminant analysis classifier (CoSMoMVPA; regularisation 0.01) was trained and tested with leave-one-reshuffle-out cross-validation (one reshuffle = one SSSD and one DDDS sequence), with training classes equalised by random sub-sampling. Performance is balanced accuracy (chance = .50). For summary statistics, accuracy was averaged in the windows in which the main effects were found (S (Odd) vs. D (Odd): 0.12–0.55 s; S (Std) vs. D (Std): 0.10–0.14 s and 0.64–0.68 s) unless another window is given, and tested against chance, or paired differences against zero, with two-sided JZS Bayesian t-tests (Cauchy prior scale √2/2; N = 20). $BF_{10} \geq 3$ is taken as evidence for an effect and $BF_{01} \geq 3$ as evidence for its absence.

### S1. Generalisation to unseen shapes and colours

**Rationale**. *Same/different* decoding could in principle rely on the visual features of specific shapes or colours rather than on the relation between the two shapes of a pair. If the neural response to some shapes or colours differed systematically from that to others, a classifier could learn to associate these item-specific responses with one relation during training and then could classify test pairs from the learned relation without the relation playing any role. To test this, the classifier was evaluated only on pairs whose shapes, or whose colour, it had never encountered during training. If decoding relied on item-specific features, it should fall to chance under these conditions; if it relied on the relation, it should remain above chance.

**Methods.** We repeated Contrast 1 of the main text (Relation Type: S (Std) vs. D (Std) and S (Odd) vs. D (Odd)) and, with leave-shapes-out cross-validation, Contrast 2 (Sequence Role: Std (S) vs. Odd (S) and Std (D) vs. Odd (D)), replacing the leave-one-reshuffle-out cross-validation with two stricter schemes in which the test pairs contained shapes, or a colour, that were absent from training. In leave-shapes-out cross-validation, each fold was defined by one set of shapes (a single shape for a *same* pair, two shapes for a *different* pair) and one held-out reshuffle: the classifier was tested on the pairs of the held-out reshuffle that contained exactly that set of shapes, and trained on the pairs of all other reshuffles that contained neither shape. A test pair therefore never shared a shape with any training pair; holding out a single shape would not ensure this, because a *different* pair would still share its other shape with the training set. Leave-colours-out cross-validation was identical, with the colour of the pair in place of its shapes. Because each fold contained pairs of only one relation type, predictions were pooled across folds for each relation type, and balanced accuracy was computed as the mean of the accuracies for *same* and *different* pairs.

**Results**. Under both stricter schemes, *same/different* decoding remained close to that obtained with the original cross-validation, in which the same shapes and colours appeared in training and test (Fig. S1). At the oddballs, S (Odd) vs. D (Odd) remained well above chance with leave-shapes-out ($M$ = .539) and with leave-colours-out cross-validation ($M$ = .544; both $BF_{10}$ > 10,000). At the second standards, the weaker S (Std) vs. D (Std) effect was also above chance ($M$ = .512–.516, $BF_{10}$ = 3.8–67). For Contrast 2, Sequence Role remained above chance with leave-shapes-out cross-validation, close to the original cross-validation (0.20–0.60 s: Std (S) vs. Odd (S) $M$ = .552, $BF_{10}$ = 170, versus .554; Std (D) vs. Odd (D) $M$ = .534, $BF_{10}$ = 18, versus .543).

**Summary.** *Same/different* decoding did not depend on specific shapes or colours. When the classifier was tested only on pairs whose shapes or whose colour it had never been seen during training, the oddballs were still decoded well above chance, and the weaker effect at the standards also survived. Decoding of Sequence Role likewise did not depend on the particular shapes presented.

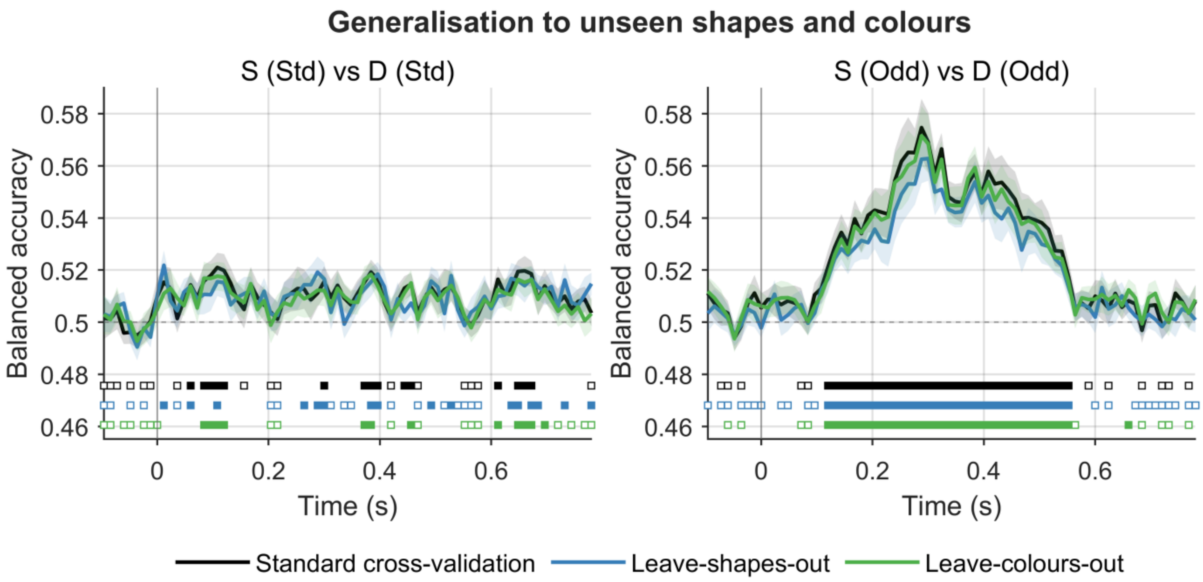


**Fig. S1.** Generalisation to unseen shapes and colours. *Same/different* decoding at the second standards (S (Std) vs. D (Std), left) and at the oddballs (S (Odd) vs. D (Odd), right). Black: original leave-one-reshuffle-out cross-validation. Blue: leave-shapes-out, in which the test pairs shared no shape with any training pair. Green: leave-colours-out, in which the test pairs shared no colour with any training pair. Lines show the group mean ($N = 20$) and shading ±1 SEM; the dashed line marks chance (.50) and the vertical line stimulus onset. Squares beneath the curves (one row per curve, in the curve's colour) mark time windows with $BF_{10} \geqslant 3$ for above-chance decoding (filled) or $BF_{01} \geqslant 3$ for chance-level decoding (open).

**S2. Stimulus familiarity**

**Rationale**. Because the set of *same* pairs was much smaller than the set of *different* pairs (180 versus 1,620 unique images), each *same* image was shown more often than each *different* image (on average 8.4 versus 1.5 times per session). Images can evoke weaker or otherwise altered responses as they become familiar, so *same* and *different* pairs could have been distinguished by their familiarity rather than by their relation. If familiarity drove decoding, decoding should decrease when *same* and *different* oddballs are matched on how often each image had been seen, and it should grow over the session, because *same* images are repeated while *different* images remain mostly novel.

**Method**. For every presentation we counted its prior exposures: the number of earlier presentations of the identical image (same shapes, colour and left–right order) in the session. (i) Exposure matching: *same* and *different* oddballs were matched for prior exposure (0, 1 or 2) within each reshuffle by randomly sub-sampling the larger class in every reshuffle × exposure cell. This equated familiarity (mean prior exposure 0.55 for both classes, compared with 1.74 for same and 0.23 for *different* oddballs in the full data) and retained about one fifth of the presentations (on average 68 per class). Because using fewer presentations lowers accuracy by itself, the matched result was compared with a count-matched reference: the same number of presentations per class and reshuffle, chosen at random without regard to exposure. Sampling was repeated 10 times and accuracies averaged. (ii) Time in session: because *same* images accumulate exposures over the session whereas *different* images remain mostly novel, a familiarity account predicts that *same/different* decoding increases over the session. To test this, accuracy in the folds of the original cross-validation that tested the first four reshuffles was compared with accuracy in the folds that tested the last four.

**Result**. With *same* and *different* oddballs matched on prior exposure, S (Odd) vs. D (Odd) decoding remained above chance ($M = .526$, $BF_{10} = 43$) and did not differ from the count-matched reference ($\Delta = +.005$, $BF_{01} = 2.8$). *Same/different* decoding did not differ between the first four and the last four reshuffles (S (Odd) vs. D (Odd): .548 early versus .544 late, $BF_{01} = 2.3$; S (Std) vs. D (Std): late − early $BF_{01} = 3.0$ and 4.3 in its two windows; Fig. S2). For S (Std) vs. D (Std), exposure matching left too few presentations for even the count-matched reference to exceed chance, so familiarity at the second standards is addressed by the comparison across the session only.

**Summary**. *Same* images were shown more often than *different* images, but this difference in familiarity does not explain *same/different* decoding at the oddballs. Decoding remained above chance when *same* and *different* oddballs were matched on how often each image had been seen before, and it did not increase over the session, at the oddballs or at the second standards, as *same* images became more familiar.

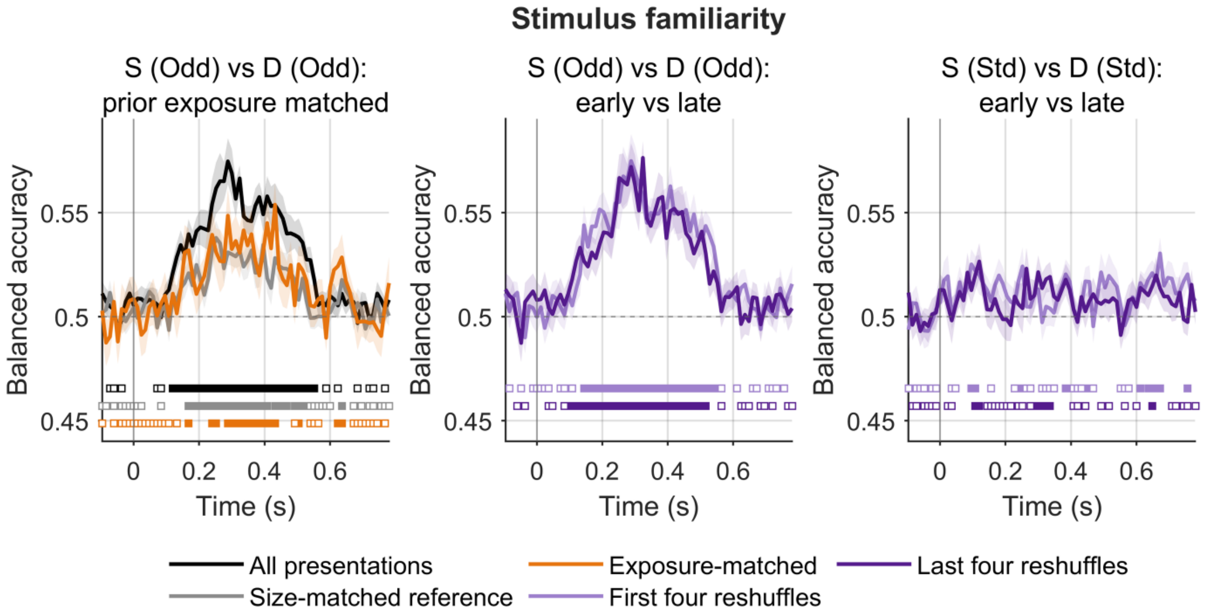


**Fig. S2.** Stimulus familiarity. Left: S (Odd) vs. D (Odd) on all presentations (black); on *same* and *different* oddballs matched on prior exposure (0–2 earlier presentations of the identical image) within each reshuffle (orange); and on a count-matched reference with the same number of presentations per class and reshuffle, sampled without regard to exposure (grey). Matched and reference curves are averages over 10 random samples. Centre and right: accuracy of the original cross-validation in the folds testing the first four (light purple) and the last four (dark purple) reshuffles, for S (Odd) vs. D (Odd) and S (Std) vs. D (Std). Lines show the group mean (N = 20) and shading ±1 SEM; the dashed line marks chance (.50) and the vertical line stimulus onset. Squares beneath the curves (one row per curve, in the curve's colour) mark time windows with $BF_{10} \geqslant 3$ for above-chance decoding (filled) or $BF_{01} \geqslant 3$ for chance-level decoding (open).

### S3. Amplitude normalisation

**Rationale**. *Same* and *different* oddballs could be separable simply because one evokes a larger response than the other. In that case, the scalp patterns would differ only in overall amplitude, and the classifier would be detecting response magnitude rather than a difference in how activity is distributed across channels and time. To test this, the overall amplitude of every pattern was removed before classification. If decoding relied on amplitude alone, it should fall to chance after this normalisation; if the two relations evoke differently shaped patterns, it should remain above chance.

**Method.** Before classification, the feature vector of every presentation in every window (64 channels × 5 samples) was *z*-scored across its 320 values: its mean was subtracted and it was scaled to unit variance. This removes the overall size and offset of the response while keeping its shape across channels and time, making the classifier insensitive to the overall amplitude of a pattern, as in correlation-based pattern analysis (Haxby et al., 2001). S (Std) vs. D (Std) and S (Odd) vs. D (Odd) were then decoded as in the main analysis.

**Result.** Decoding was almost unchanged (0.12–0.40 s: S (Odd) vs. D (Odd) M = .542, $BF_{10}$ = 7676, versus .549 on raw patterns; S (Std) vs. D (Std) M = .507, $BF_{10}$ = 4.1; Fig. S3). *Same* and *different* therefore differ in the shape of the scalp pattern, not only in its size.

**Summary.** *Same* and *different* were not distinguished merely because one evoked a larger response. After the overall size of every activity pattern was removed, decoding was almost unchanged, so the two relations evoked differently shaped scalp patterns.

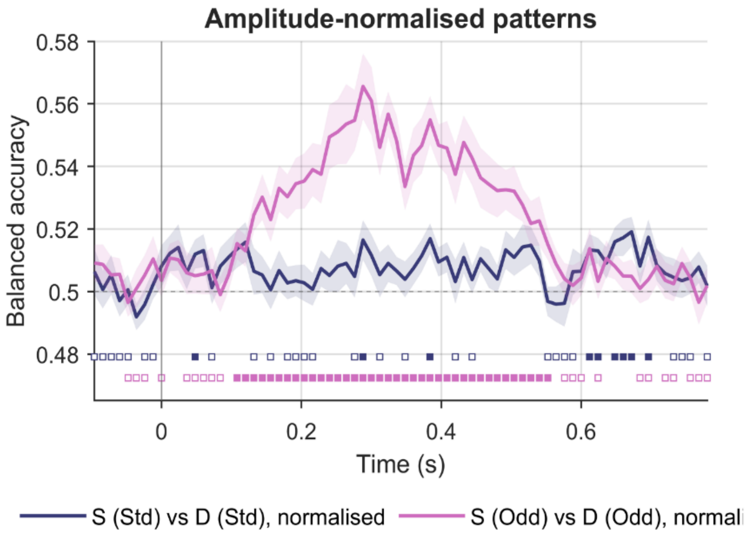


**Fig. S3.** Amplitude normalisation. *Same–different* decoding at the second standards (S (Std) vs. D (Std), indigo) and at the oddballs (S (Odd) vs. D (Odd), magenta) after the activity pattern of every presentation in every 20-

ms window (64 channels × 5 samples) was z-scored, removing overall response size and offset. Lines show the group mean (N = 20) and shading ±1 SEM; the dashed line marks chance (.50) and the vertical line stimulus onset. Squares beneath the curves (one row per curve, in the curve's colour) mark time windows with $BF_{10} \geqslant 3$ for above-chance decoding (filled) or $BF_{01} \geqslant 3$ for chance-level decoding (open).

### S4. Restricting decoding to posterior channels only

**Rationale**. No electro-oculogram was recorded, so eye movements and blinks could not be removed from the data used for decoding. Ocular signals are largest over frontal electrodes, but through the average reference they spread to every channel, so a classifier could in principle separate conditions using ocular activity that differed between them. For example, if participants blinked more after one kind of oddball. Restricting decoding to posterior channels re-referenced to their own mean removes this route. If the effects were carried by ocular activity, they should be reduced or absent when decoding is restricted to these channels.

**Method.** All decoding contrasts (standard vs. oddball within each sequence, and Contrasts 1 and 2 of the main text) were decoded again from 20 posterior channels (P1–P10, Pz, PO3, PO4, PO7, PO8, POz, O1, O2, Oz and Iz) re-referenced to their own mean. Re-referencing within this set removes the average reference, so frontal and ocular activity cannot reach these channels through it. Features were the 20 channels × 5 samples of each window; all other settings were as in the main analysis. Accuracy from the posterior set was compared with accuracy from all 64 channels with the average reference.

**Result.** Posterior channels reproduced every contrast with the same size and time course (0.12–0.40 s: S (Odd) vs. D (Odd) .556 versus .549 from all channels; S (Std) vs. D (Std) .512 versus .511; all-channel − posterior differences $\leqslant$ .008 in every contrast, $BF_{01}$ up to 4.3; Fig. S4).

**Summary.** Eye movements and blinks do not explain the results. Decoding from posterior channels only, which ocular activity cannot reach through the reference, reproduced every main-text contrast with the same size and time course.

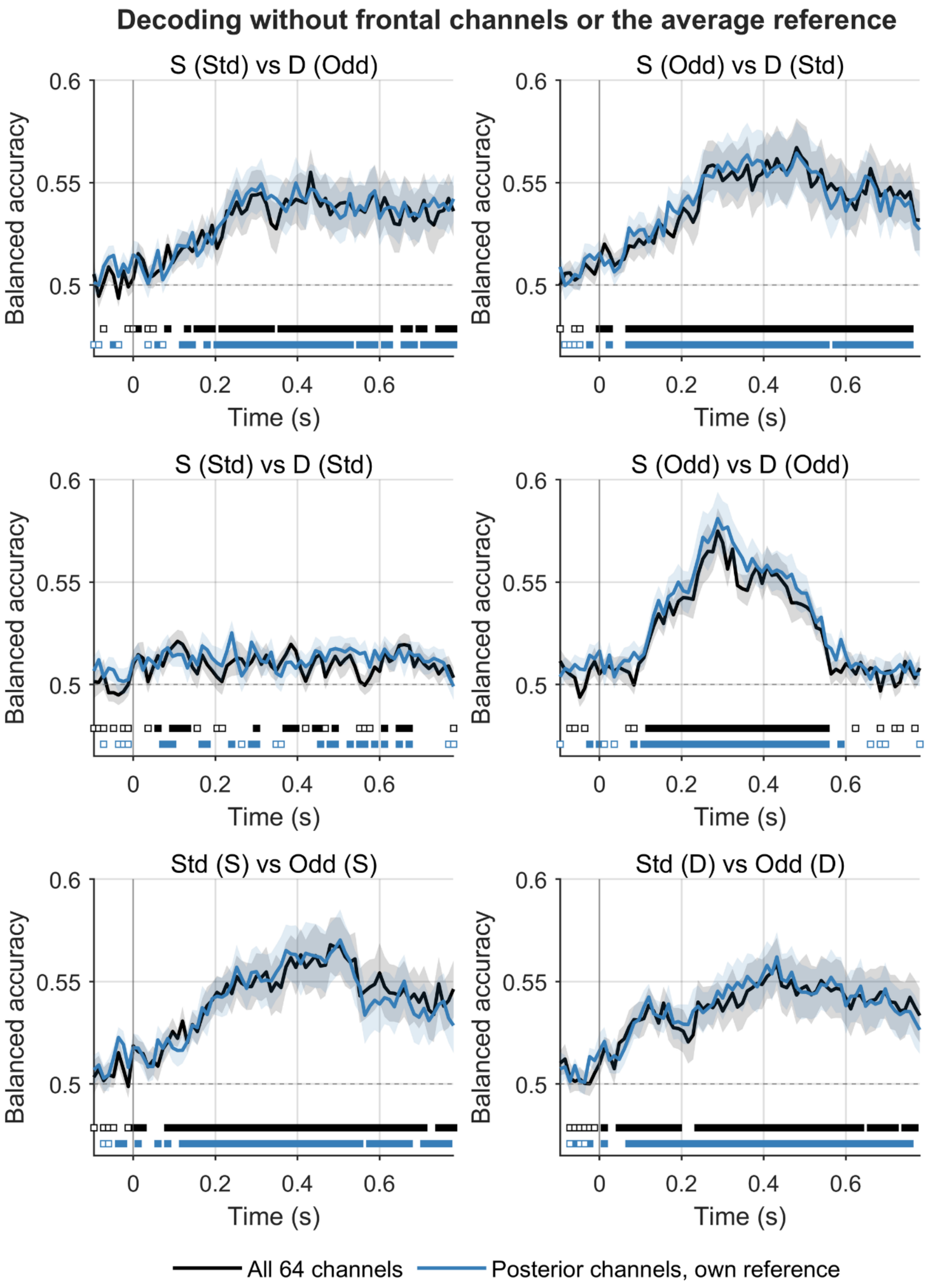


**Fig. S4. Restricting decoding to posterior channels only. Decoding of all contrasts (rows: standard vs. oddball within each sequence; Contrast 1, Relation Type; Contrast 2, Sequence Role)** from all 64 channels with the average reference (black) and from 20 posterior channels (P, PO, O and Iz) re-referenced to their own mean, which ocular activity cannot reach through the reference (blue). Lines show the group mean (N = 20) and shading ±1 SEM; the dashed line marks chance (.50) and the vertical line stimulus onset. Squares beneath the curves (one row per curve, in the curve's colour) mark time windows with $BF_{10} \geqslant 3$ for above-chance decoding (filled) or $BF_{01} \geqslant 3$ for chance-level decoding (open).
